# Evaluating Complexity of Fetal MEG Signals: A Comparison of Different Metrics and Their Applicability

**DOI:** 10.1101/540278

**Authors:** Julia Moser, Siouar Bensaid, Eleni Kroupi, Franziska Schleger, Fabrice Wendling, Giulio Ruffini, Hubert Preißl

## Abstract

In this work, we aim to investigate whether information based metrics of neural activity are a useful tool for the search for consciousness before and shortly after birth. Neural activity is measured using fetal magnetoencephalography (fMEG) in human fetuses and neonates. Based on recent theories on consciousness, information-based metrics are established to measure brain complexity and to assess different levels of consciousness. Different metrics (measures of entropy, compressibility and fractality) are, thus, explored in a reference population and their usability is evaluated. For comparative analysis, two fMEG channels were selected: one where brain activity was previously detected and one at least 15cm away, that represented a control channel. The usability of each metric was evaluated and results from the brain and control channel were compared. Concerning the ease of use with fMEG data, Lempel-Ziv-Welch (LZW) compression was evaluated as best, as it is unequivocal and needs low computational effort. The fractality measures have a high parameter space and therefore forfeit comparability, while entropy measures require a higher computational effort and more parameters to adjust compared to LZW. Comparison of a channel with brain activity and a control channel in neonatal recordings showed significant differences in most complexity metrics. This clear difference can be seen as proof of concept for the usability of complexity metrics in fMEG. For fetal data, this comparison produced less clear results which can be related to leftover maternal signals included in the control channel. Further work is necessary to conclusively interpret results from the analysis of fetal recordings. Yet this study shows that complexity metrics can be used for fMEG data on early consciousness and the evaluation gives a guidance for future work. The inconsistency of results from different metrics highlights the challenges of working with complexity metrics as neural correlates of consciousness, as well as the caution one should apply to interpret them.

## 1 Introduction

Consciousness is known to be one of the characteristics that make humans unique. But when does this aspect of the human mind arise? Is it possible that consciousness already exists before birth? From the 24^th^ week of gestation, a fetus can process sensory stimuli at a cortical level, as thalamocortical connections are already established (Kostović and Judaš, 2010). Long range pyramidal neurons – which are known to be important for conscious processing (Dehaene et al., 1998) – are developed around week 26 (Lagercrantz and Changeux, 2009). Yet, it is difficult to assess the conscious state of a fetus in the mother’s womb. Fetal magnetoencephalography (fMEG) is a tool to noninvasively investigate fetal brain activity in the last trimester of pregnancy and in neonates shortly after birth (Preissl et al., 2004). This tool makes it possible to measure a neural correlate of consciousness in fetuses and pursue the question of the debut of consciousness in human life.

During the last decades, work in the field of disorders of consciousness led to an increased interest in neural correlates of consciousness. One way to find such a correlate is to look at complexity of neurological data (Tononi and Edelman, 1998). In nature, complexity of physiology is related to the adaptive capacity of an organism (Costa et al., 2002). This is translated into physiological signals with long-range correlations across various spatio-temporal scales – a behavior that is named self-organization – that indicate the presence of self-invariant and self-similar structures (Pritchard and Duke, 1995). The self-organizational properties of a complex system can be quantified by estimating its dimension (Theiler, 1990), or its ability to compress information (Cover and Thomas, 2012; Ruffini, 2017a).

For a system to be complex, it has to operate on several scales and also show an interplay between those scales (Lutzenberger et al., 1995). This is a property of a so-called chaotic system, and can be measured in space and time (Elbert et al., 1994). Typically chaos in space is estimated with the fractal dimension, which is defined as the dimension of a strange attractor towards which a complex system evolves in phase-space (Grassberger and Procaccia, 1983b). Thus, the fractal dimension, namely D, describes the overall complexity of an object, which can be the geometrical complexity, the space filling property, the roughness of a surface, or the variation of a time series. The fractal dimension D is defined by the logarithmic ratio of change in detail with change in scale (Di Ieva, 2016). This relates to the distinct characteristic of a fractal, namely the property of self-similarity: i.e., pieces of an object are similar to larger pieces of it as well as to the whole object (Eke et al., 2002). In nature, fractals are usually only statistically self-similar which means that smaller excerpts are not necessarily exact copies of the larger ones, but they are the same on average (Pritchard and Duke, 1995). In contrast to fractal dimension, entropy gives information about the dynamical properties of an attractor and not about its geometrical shape (Rodríguez-Bermúdez and Garcia-Laencina, 2015). Chaos in time therefore relates to this stability and sensitivity to initial conditions (Elbert et al., 1994; Baranger, 2000). Related to those properties of complex systems, several measures were developed to quantify their complexity. The sensitivity to initial conditions can be quantified in terms of the Lyapunov exponent and the Kolmogorov entropy, also known as information dimension (Theiler, 1990). Different entropy measures as well as the measure of compressibility can be employed for this. Ruffini (2017a) recently proposed a theory of consciousness that considers the brain an engine that strives to model the world with simplicity, while learning it is a result of exchanging information with it. According to this theory, the ability of the brain to compress information is an indicator of consciousness.

In consciousness neuroscience, this quantification of complexity is used in different scenarios. The main areas of application are research with patients with disorders of consciousness, anesthesia monitoring and sleep studies (e.g. Casarotto et al., 2016). For instance, Burioka et al. (2005) showed that approximate entropy calculated on small segments of electroencephalography (EEG) data decreases with depth of sleep. In particular, there is a linear decrease from wakefulness over sleep stage one until four, while rapid eye movement sleep showed values similar to wakefulness. Analysis of data from EEG, MEG and intracranial EEG recordings confirmed this drop of complexity with a drop in wakefulness (Mateos et al., 2018). Complexity was calculated with entropy and compressibility measures. Zhang et al. (2009) could differentiate between active sleep and quiet sleep in newborns by means of sample entropy. Similarly, the dimensional complexity of the EEG pattern measured by correlation dimension (CD) was found to be higher in active sleep compared to quiet sleep for infants in their first months of life (Janjarasjitt et al., 2008b; Scher et al., 2009). Furthermore, scale free properties caused by the self-similarity of fractals, can be used to differentiate between sleep stages whereas an increase or decrease of values depends on the scale free parameter estimated (Weiss et al., 2009).

In general, the measurement of this scale free behavior appears promising in the investigation of state transitions (Weiss et al., 2009). Studies with Propofol anesthesia showed a change in scale free behavior before and after loss of consciousness (Eagleman et al., 2018) and a difference between wakefulness and loss of consciousness as well as recovery from anesthesia (Tagliazucchi et al., 2016). Also with the help of entropy measures, Eagleman et al. (2018) could show a change in complexity of scalp EEG data before and after loss of responsiveness in anesthesia patients. Similarly, Schartner et al. (2015) could distinguish between loss of consciousness during Propofol induced anesthesia and wakeful rest by means of entropy measures as well as compressibility measures. Furthermore, higher entropy values were shown in the EEG data of healthy control participants compared to unresponsive wakefulness patients, matched for sex and age (Sarà and Pistoia, 2010).

Compressibility measures in combination with transcranial magnetic stimulation are widely used to distinguish between patients with different disorders of consciousness (e.g. unresponsive wakefulness state, minimally conscious state, locked in syndrome) and healthy subjects in different sleep stages. In addition, sedation with different anesthetics can be differentiated (e.g. Propofol and Ketamine), whereby in both cases patients are behaviorally unresponsive but in case of Ketamine they report vivid dreams (e.g. Casali et al., 2013; Sarasso et al., 2015; Casarotto et al., 2016; Bodart et al., 2017; Rosanova et al., 2018). Moreover, a recent study on MEG revealed increased LZW (Lempel-Ziv Welch) complexity during a psychedelic state of consciousness induced using Ketamine, LSD, and Psilocybin compared to a placebo effect (Schartner et al., 2017). Regarding other, related, patient populations, schizophrenic patients have higher LZW complexity compared to healthy controls (Fernández et al., 2011), and depressed patients have higher MEG pre-treatment complexity that decreases after 6 months of pharmacological treatment (Méndez et al., 2012).

These findings show that there are many valid approaches to use complexity metrics to quantify consciousness, although to our knowledge there are no studies that investigated complexity on early consciousness. Yet, the exact relation of the different metrics is hard to grasp and it is difficult to define a metric that is most suitable for a specific purpose. Therefore, in the current study, as we use a different type of data than previous studies, we follow an explorative approach and apply numerous different metrics to our fMEG data (Table 1). The aim of this approach is to capture different aspects of complexity and compare the metrics regarding their behavior towards this special type of data and their usability when employing them to pursue a novel question. In particular, data from fetal MEG recordings with different gestational ages and additional neonatal recordings are used in this study. The data included in the analysis previously showed auditory event-related responses, which allowed for identification of channels with high brain activity within the sensor space. Such data is used, as it is otherwise difficult to localize clusters of brain activity. The goal of the analysis is to evaluate if complexity metrics are a useful tool for fMEG analysis in the search for fetal consciousness.

**Table 1:** Overview used Metrics

| Concept investigated | Method | Objective |
| --- | --- | --- |
| **Entropy** | Multiscale Entropy | Measurement of self-similarity of time series by looking at repeating sequences on multiple scales |
| **Entropy** | Multiscale Permutation Entropy | Measurement of self-similarity of time series by looking at probability of patterns in data on multiple scales |
| **Compressibility** | Lempel-Ziv-Welch compression | Quantification of compressibility of time series |
| **Fractality** | Correlation Dimension | Measurement of strangeness of attractor, towards which complex system evolves |
| **Fractality** | Scale free approaches | Detection of power-law exponent that describes scale free behavior and additionally description of multifractal properties |

**Table 2:** Comparison of usability of methods

| Method | Computational Effort | Parameter Space | Comparability | Overall Usability |
| --- | --- | --- | --- | --- |
| MSE | moderate | moderate | high | OK |
| MPE | moderate | moderate | moderate | OK |
| LZW | very low | small | high | good |
| CD | high | large | moderate | weak |
| MF DFA | low | large | low | weak |
| WLB MF | low | large | low | weak |

## 2 Material and Methods

### 2.1 Fetal Magnetoencephalography

fMEG is a non-invasive tool to measure heart and brain activity in fetuses in the last trimester of pregnancy and in neonates shortly after birth (Preissl et al., 2004). For the recording of fetal- and neonatal data, the SARA (SQUID Array for Reproductive Assessment, VSM MedTech Ltd., Port Coquitlam, Canada) system installed at the fMEG Center at the University of Tübingen is used (Figure 1). To attenuate magnetic activity from the environment, the device is installed in a magnetically shielded room (Vakuumschmelze, Hanau, Germany). The system includes 156 primary magnetic sensors and 29 reference sensors. The magnetic sensors are distributed over a concave array whose shape is designed to match the maternal abdomen. Based on an ultrasound measurement (Ultrasound Logiq 500MD, GE, UK) prior to the fMEG recording the position of the fetal head is determined and is marked by a localization coil placed on the maternal abdomen. Three additional localization coils are placed on the spine, left and right side of the subject, to track position changes in relation to the sensor array. An ultrasound directly after the measurement is used to confirm the fetal head position. In case of a change in position, datasets are excluded. Auditory stimulation can be presented via a balloon placed between the maternal abdomen and the sensor array. Neonates get a small, child-appropriate earphone (Ear Muffins from Natus, Biologic, San Carlos, USA), placed on one ear. For neonatal recordings a cradle is attached to the fMEG device. The neonate is attended by one parent inside the measurement room and is measured asleep or quiet awake.

**Figure 1:**
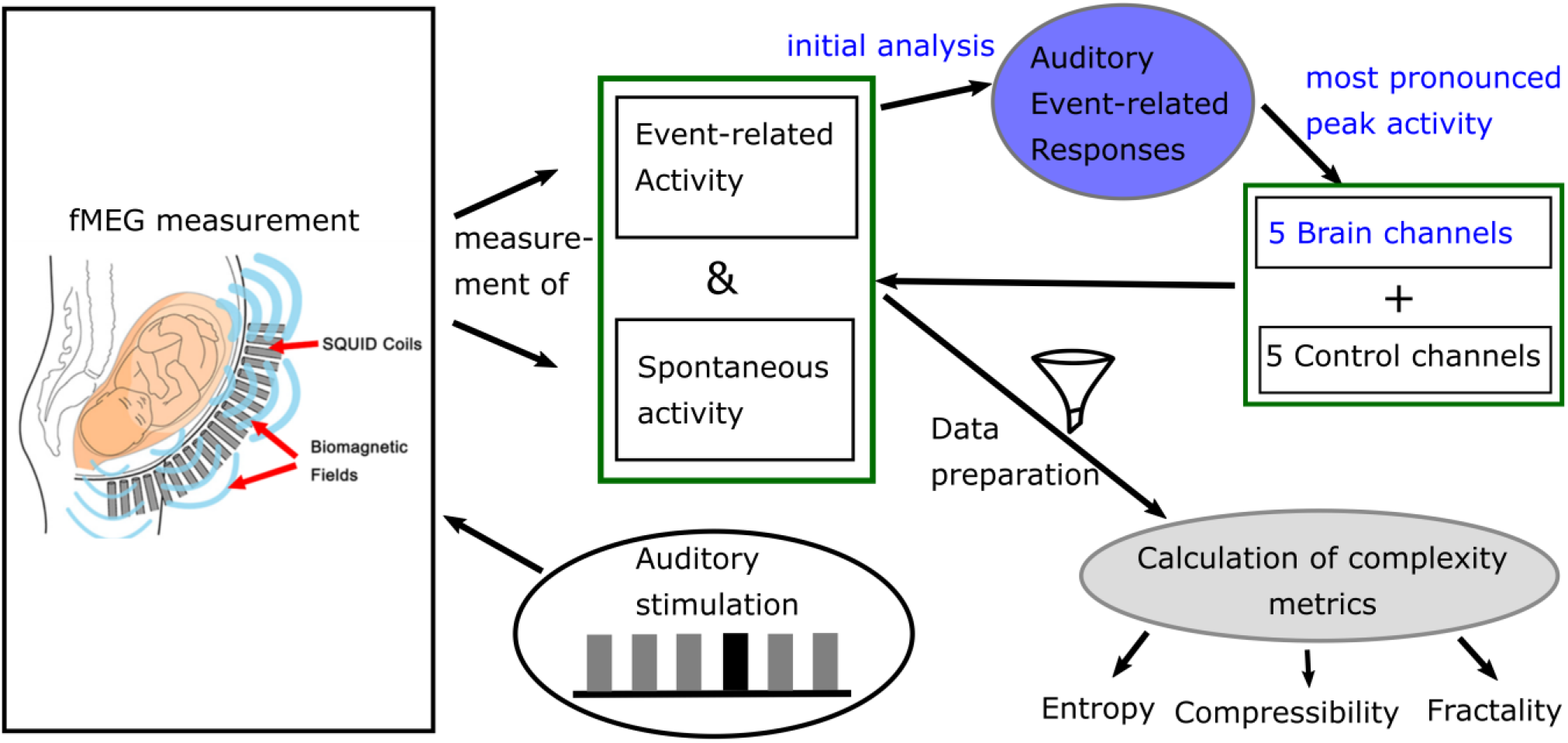
Diagram of data acquisition and processing. Blue: analysis done for previous studies.

### 2.2 Dataset

For the current analysis, fetal and neonatal data from previously analyzed studies was used (Linder et al., 2014; Morin et al., 2015). The auditory stimulation paradigm used in these studies consists of an auditory oddball paradigm with a 500Hz tone as standard and a 750Hz tone as deviant. The standard occurs in 80% of times in a pseudorandomized order. Each tone is 500ms long with an inter-trial interval of 1500ms. 45 fetal recordings were selected from subjects where auditory event-related responses were detected. They have a gestational age range from 29 to 39 weeks – 15 of them in an early phase of the third trimester (29-32 weeks), 15 in the middle (33-36 weeks) and 15 in a later phase (37-39 weeks) – approximately uniformly distributed over the whole age range. 15 neonatal recordings were included with an age range from 4 to 46 days (mean=17.47; SD=12.68). For all subjects, data with auditory stimulation (“audio”) and data without stimulation (“spont”) is available. As the length of the fetal datasets varies from six to 15 minutes, for all of them only the first six minutes were used. The neonatal datasets all have a length of ten minutes.

### 2.3 Data Analysis

#### 2.3.1 Preprocessing

As first step for all datasets, the maternal magnetocardiogram and fetal magnetocardiogram were detected by template matching or using the Hilbert transform algorithm and were subtracted from the relevant signal, through signal space projection (Vrba et al., 2004; McCubbin et al., 2006; Wilson et al., 2008). One of the two methods was selected depending on which method yielded better results, which is the established procedure for fetal brain analysis (e.g., Linder et al., 2014). Matlab 16b was used (The MathWorks, Natick, MA) for all further processing steps, expect for the calculation of the compressibility measure, where Python 2.7 was used (Python Software Foundation, https://www.python.org/). Fieldtrip (Oostenveld et al., 2010) was used to filter all fetal data between 0.5 and 10Hz and all neonatal data between 0.5 and 15Hz which is the usual filtering range for the analysis of evoked brain responses (Linder et al., 2014; Schleger et al., 2014). Additionally, data was down sampled from 610.35Hz to 256Hz.

In the previous analysis of these datasets, five channels were selected for the analysis of auditory event-related responses (example for analysis procedure in Schleger et al., 2014). As these were the five channels with the highest evoked brain activity, they were selected as brain channels for this study (“brain”). Additional to those five channels, five control channels (“control”) were selected that are all more than 15cm away from the brain channels. By default, the five channels were situated in the upper middle part of the sensor field. However, if the fetus was positioned in a way that was too close to those sensors, five channels in the lower middle part were selected. As the five brain channels showed similar behavior, for simplicity of the later analysis only the brain channel with the highest amplitude, and one control channel, were used. The usability of a single channel for complexity analysis was previously demonstrated (e.g., Scher et al., 2005). For later analysis the signal was cut into windows of 1 minute (Scher et al., 2009; Kaffashi et al., 2013), to generate a more stationary signal. This results in six time-windows for fetal data and ten time-windows for neonatal data. If a time window included a signal that was higher than 1pT, it was classified as containing an artefact and the concerning time window was removed from the analysis.

#### 2.3.2 Complexity Metrics

To measure the informational complexity of the fMEG signal, multiscale entropy (MSE) and multiscale permutation entropy (MPE) was used, as well as LZW compression. Additionally, we included the geometrical properties of the signal and measured its amount of fractality. As an approximation for this, the CD, also known as dimensional complexity (Janjarasjitt et al., 2008b), is calculated. In addition, scale free behavior, which is a basic property of a fractal, was taken into account.

#### 2.3.3 Multiscale Entropy

MSE calculates sample entropy (SE) for different time scales. If SE is calculated, lower values indicate more self-similarity in time series and the calculation is largely independent of recording length and relatively consistent (Richman and Moorman, 2000). This traditional entropy measure takes only one scale into account, therefore MSE uses a coarse-grained time series with different scaling factors which takes long range correlations into account (Costa et al., 2002). By considering multiple scales, both highly deterministic and completely random signals result in low values, only complex signals can reach a high value (McIntosh et al., 2008).

For the coarse graining step, the dataset is divided into non-overlapping windows of size τ and the data points in each window are averaged. SE is based on the definition of Kolmogorov entropy and is defined as the negative logarithm of the probability of two sequences that are similar for m points, to be similar at the point m+1 as well (Richman and Moorman, 2000). r describes the tolerance within which two points are accounted as similar (Richman & Moorman, 2000).

For the calculation of MSE the “msentropy” function out of the WFTB Toolbox (Goldberger et al., 2000; Silva and Moody, 2014) was used. The parameters used in the current analysis are oriented at the default parameters described by Costa et al. (2005). We used m = 2, r = 0.15 (15% of the standard deviation of the time series) and N = 15360 which is a bit lower than the recommended N = 2 * 10^4 but was selected in terms of comparability of the different methods and did not show any disadvantage compared to an example analysis with a larger N. We chose τ to contain all values between 1 and 20 to calculate the SE for 20 different scales whereas scale one equals the original time series. A time series is considered as more complex than another if a majority of scales show higher entropy values (Costa et al., 2005). For that reason we used the average MSE over all scales for all further comparisons.

#### 2.3.4 Multiscale permutation Entropy

MPE calculates permutation entropy (PE) for multiple time scales. In comparison to other complexity measures PE is very robust towards noise (Bandt and Pompe, 2002; Zanin et al., 2012). PE looks at different patterns within a time series with the idea that those patterns do not have the same probability of occurrence and that this probability can be informative regarding the underlying dynamics of the system (Zanin et al., 2012). PE uses short samples of a time series to look at their permutation patterns and their frequency of occurrence in relation to all possible permutation patterns (Bandt and Pompe, 2002). PE can be used to quantify complexity of a dynamical time series as it refers to its local order structure (Ouyang et al., 2013). A large value of PE indicates that all permutations are equally likely, a value close to zero signifies a very regular time series (Ouyang et al., 2013).

MPE was calculated using the “MPerm” function by Ouyang (2012, November 21)^1^. Like for MSE, the first step is a coarse graining where we selected the same τ as in the MSE calculation which is denoted as s in the MPE analysis. In this study m = 4 was selected for the length of the short samples and a time delay t = 1 which corresponds to the default value. Like for the MSE, the average MPE over all scales was used for further comparisons.

#### 2.3.5 Lempel-Ziv-Welch Compressibility

The LZW compression is a measure closely related to Kolmogorov complexity and Shannon entropy (Gao et al., 2011), and is originally described by Ziv and Lempel (1978). For LZW compression, a dictionary that starts with the shortest new sequence in a time series is built and then adds longer sequences until it captures all non-repetitive sequences (Ruffini, 2017b). The length of this dictionary defines the amount of compressibility of a time series (Aboy et al., 2006). LZW values increase with increasing frequency but not with increasing amplitude as well as with increasing power of noise and increasing signal bandwidth (Aboy et al., 2006).

To calculate LZW, a signal of length n has to be binarized, which in our case is done by a median split as a threshold. In particular, values below the median are indicated as zero, whereas values above the median as one. The median split is relatively robust to outliers compared to other methods (Aboy et al., 2006). After the binarization process, the data can be compressed to a set of “words”, c(n), and the description length of the dictionary is defined as the number of included words times the bits needed to encode those words plus the bits needed to define a new symbol in the dictionary (Ruffini, 2017b).

The complexity counter, c(n), is then normalized by the length of the data string (Aboy et al., 2006). In the current analysis, ρ0, which represents the c(n) normalized by the original string length, is the value used to indicate LZW compressibility. A higher ρ0 value indicates higher complexity, thus, less ability to compress. For a more detailed description of the process of compression, the reader is referred to Aboy et al. (2006), and for the algorithm used in the current analysis to Ruffini (2017b).

#### 2.3.6 Correlation Dimension

The CD is a measure for the strangeness of an attractor, which is closely related to the fractal dimension D. In addition to the geometrical properties of the attractor it takes the dynamics of coverage of the attractor into account (Grassberger and Procaccia, 1983b). The CD uses the statistics of pairwise distances to estimate dimension and is based on the scaling of mass with size (Theiler, 1990). Correlations between points of long-time series on the attractor are used for that (Grassberger and Procaccia, 1983b). For a complete description of a higher dimensional nonlinear system, a time series – which is an observation in one dimension – has to be unfolded into a higher dimensional space, the so called “embedding space” (Janjarasjitt et al., 2008a). The process of embedding a time series in a higher dimension is described by (Takens, 1981).

The CD is calculated using the correlation integral, defined as the ratio of the distances between any two points that are smaller than a radius r and all possible distances (Theiler, 1990). To determine the distance in the time series of the points to be correlated, the time delay τ is introduced. τ can be set with the help of the autocorrelation function (Janjarasjitt et al., 2008a).

For the current analysis, the “gencorint” function was used to determine the CD (Grassberger and Procaccia (1983a), Albano et al. (1988), Theiler (1986) out of the Chaotic systems Toolbox, Leontitis, 2004^2^). For setting the embedding parameter, we used the false nearest neighbor algorithm (Kizilkaya (2012)^3^, Kennel et al. (1992)), and determined an embedding dimension of m = 4 for the current dataset. To determine the right time delay τ, an autocorrelation function was calculated for each subject and time window and the zero point of it was used as individual value of τ. If a time window showed an for this type of data unrealistic autocorrelation function (zero point <10), it was excluded from further analysis. Except for those two parameters all other parameters were set to the default values suggested in the function. For algorithmic efficiency, the slope of the CD can be used as an approximation of the CD (Theiler, 1990). In the current analysis, a linear fit over all points was performed to determine this slope. The slope value was used for all further comparisons.

#### 2.3.7 Scale Free Approaches

The self-similarity of a fractal can be expressed by a mathematical power-law with a distinct exponent (Eke et al., 2002). One of these exponents is the Hurst exponent (H), which is also referred to as “self-similarity parameter” (Zilber, 2014). It expresses the probability that an event is followed by a similar event and is related to the fractal dimension (D) by D = 2 – H. A value of H = 0.5 shows that a time series is uncorrelated (e.g. white noise). 0.5 < H is an indication for long range correlations and H < 0.5 for long-range anti-correlations (Kantelhardt et al., 2002). For some fractal processes one power-law exponent is not enough to characterize them and a multifractal formalism can be used to describe several exponents (Di Ieva, 2016). In case of multifractality, the scaling function of a signal is not linear anymore and can thus not be described by a single scaling exponent H but by a nonlinear scaling function (Zilber, 2014). Those multiple H are called Hölder exponents and can be used to span a multifractal spectrum. The width of this spectrum (M) is a measurement for the amount of multifractality (Zilber, 2014).

Self-similar processes also described as 1/*f* or scale free behavior can be mainly observed in the infraslow frequency range of the power spectrum (Zilber, 2014). This knowledge is important for the selection of scale ranges in scale free analysis. For the following analysis, the focus is on self-similar processes in the range 0.5-2 Hz. To assess H and M, two methods for multifractal analysis are employed. For both, all data was normalized to avoid the influence of amplitude differences.

##### 2.3.7.1 Multifractal detrended fluctuation analysis

The multifractal detrended fluctuation analysis (MFDFA) is a robust analysis for the estimation of the multifractal spectrum of power-law exponents of a natural time series (Ihlen, 2012). Its basis is the detrended fluctuation analysis (DFA), one of the most popular methods to estimate scale free behavior in physiological signals which follows the idea that fluctuations within a signal are following a power-law as a function of the number of sample points (Zilber, 2014). The root mean square (RMS) of a signal is calculated over different scales (s) with n segments and a certain number of points within each segment. Larger scales are more affected by slower fluctuations, smaller scales more by faster fluctuations (Ihlen, 2012). For each scale the RMS of the individual segment (RMS(s)) – local fluctuation – is calculated, and an overall RMS (F) is computed from these values. The slope of the values of F over different scales equals H. In case of MFDFA this calculation is done for multiple orders q. As a result MFDFA obtains the set of weighted F values whose slopes obtain multiple q-order Hurst exponents. They can be used, to trace back the multifractal spectrum and its width M (Ihlen, 2012). For a more detailed description of the calculation steps, see Ihlen (2012).

The MFDFA toolbox (Ihlen (2012), MFDFA1 algorithm) was used in this study. As the algorithm is built for random-walk like signals – which are integrals of noise-like signals (Zilber, 2014) – first we checked whether the data resembles noise or random-walk signals. This is determined by the value of H, which was calculated by a simple DFA. As a cutoff H = 1.2 was selected (< noise like, > random walk like; after Ihlen, 2012). If the signal is noise like, it is transformed into a random-walk signal, and if it is random walk like, this step is skipped. As a second step, a general linear detrending of the signal is performed. For the current analysis 19 equally spaced scales ranging from 128 to 512 data points, within s, were chosen. q was selected to range from −5 to 5 in steps of 0.1 (number of q: nq = 100). The value of Hq at q = 2 equals H (Ihlen, 2012). The scale free parameters H and M were evaluated in the further analysis.

##### 2.3.7.2 Wavelet-leader based multifractal formalism

Wavelet-leader based multifractal formalism (WLBMF) poses a fast, theoretically efficient and robust analysis method for multifractal properties of real-world data (Zilber, 2014). It uses wavelet leaders to derive the multifractal properties of a signal by the knowledge of the scaling exponents.

This can be done because wavelet leaders precisely reproduce the Hölder exponents of a signal (Wendt et al., 2007). Wavelet leaders are defined as the maximum wavelet coefficients within a predefined segment (Ciuciu et al., 2008). The log-cumulants (c1-c3) of the scaling exponents give information about the shape of the multifractal spectrum. Whereas c1 equals its maximum, c2 its width and c3 its asymmetry (Wendt et al., 2007; Zilber, 2014).

For the calculation of the wavelet-leader based multifractal formalism (WLBMF) the toolbox described in Wendt et al. (2007) was used. A Daubechies wavelet with three vanishing moments was selected as a mother wavelet. The scales for the WLBMF were chosen in accordance with the scales of the MFDFA with jmin = 7 and jmax = 9. To increase the reliability of the results, Wendt et al. (2007) implemented a bootstrapping process to obtain sets of log-cumulants, which opens up new possibilities for statistical testing. In the present analysis, we used 100 bootstraps and then averaged over the bootstrapped values to retrieve the variables of interest. For a more detailed description see Wendt et al. (2007). In the present study we only evaluated c1 and c2, whereby c1 is supposed to be equivalent to H and c2 to M (Zilber, 2014).

#### 2.3.8 Statistical Analysis

For statistical analysis the results of all time windows of a certain subject and condition were averaged. Those mean values were then tested for normality with a Kolmogorov-Smirnoff test. As they showed to be not normally distributed, groups were compared with a Wilcoxon signed rank test. Significance level was set to α = 0.05 in all cases. Due to the explorative nature of the analysis no Bonferroni correction was done.

## 3 Results

### 3.1 Usability of Methods

After testing several methods for determining complexity of fMEG data, a first step was to evaluate those methods for potential future use. As shown in Table 1 we evaluated them in terms of computational costs, parameter space, comparability regarding their prevalence in the literature and summarized this with the term ‘overall usability’. Thereby a low computational effort is favored, as well as a small number of parameters that can be adjusted. Abundance of literature is seen as an advantage, to ensure comparability of results. Overall LZW had the best overall usability as it is unequivocal and fast and is therefore preferable for future analysis. Nevertheless, this evaluation concerns the present field and has only limited significance for other research questions and other types of data.

### 3.2 Detection of Brain Activity

#### Neonatal data

In the neonatal datasets, the comparison of brain vs. control resulted in a significant difference for both audio and spontaneous data for LZW, MSE, MPE and CD (p < 0.001; for MSE: p=0.016 & p=0.007). In all cases, the control channel showed higher values compared to the brain channel. For the scale free metrics in both cases H/c1 could not differentiate between brain and control but M/c2 showed a significant difference (p < 0.001). In case of M, the channel with the brain activity appeared more multifractal than the control channel, whereas in case of c2, the opposite was observed. For detailed results, see Figures 2 and 3.

**Figure 2:**
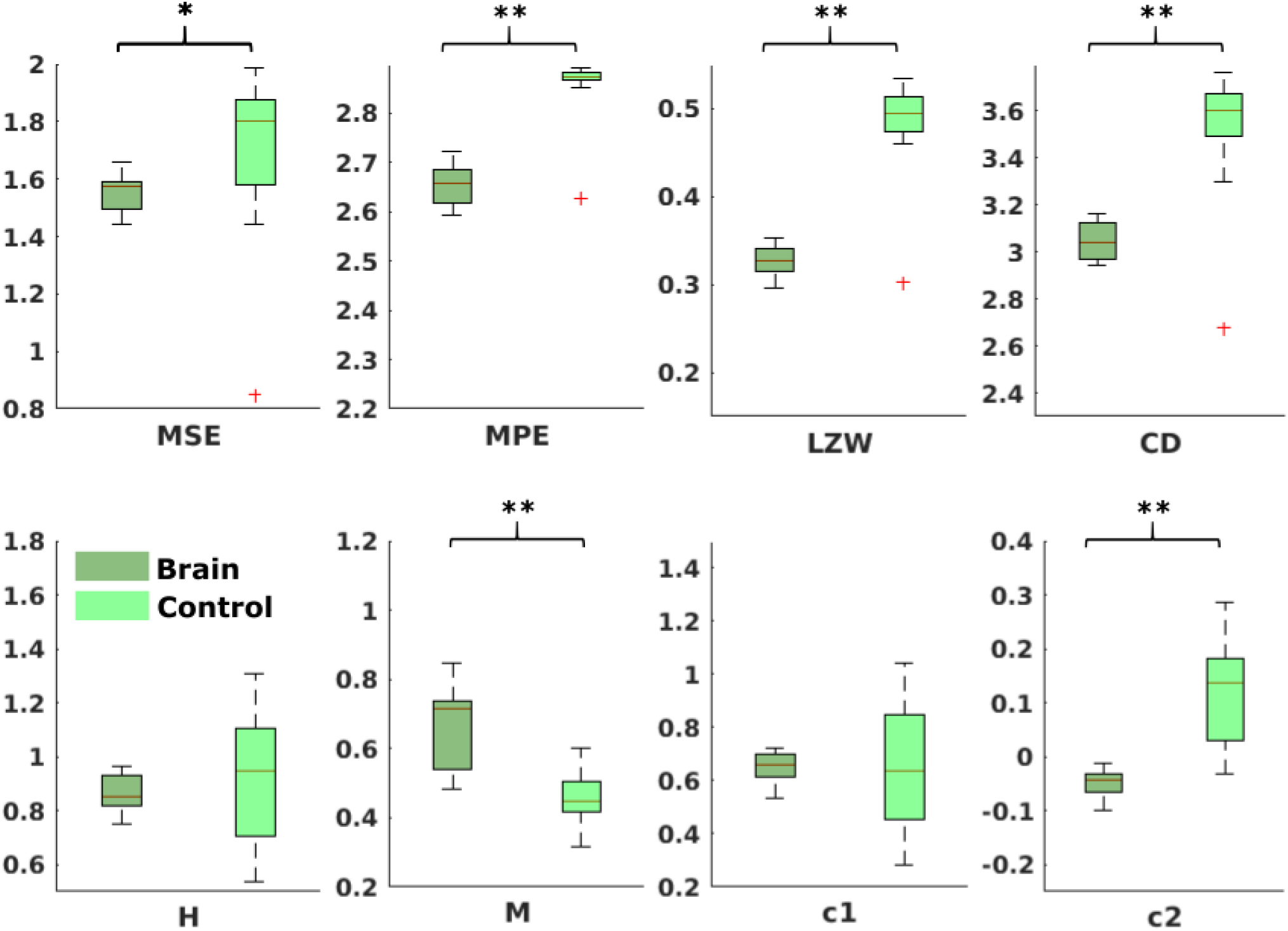
Results from analysis of different metrics on data from neonates with auditory stimulation. * depicts significant, ** highly significant, difference.

**Figure 3:**
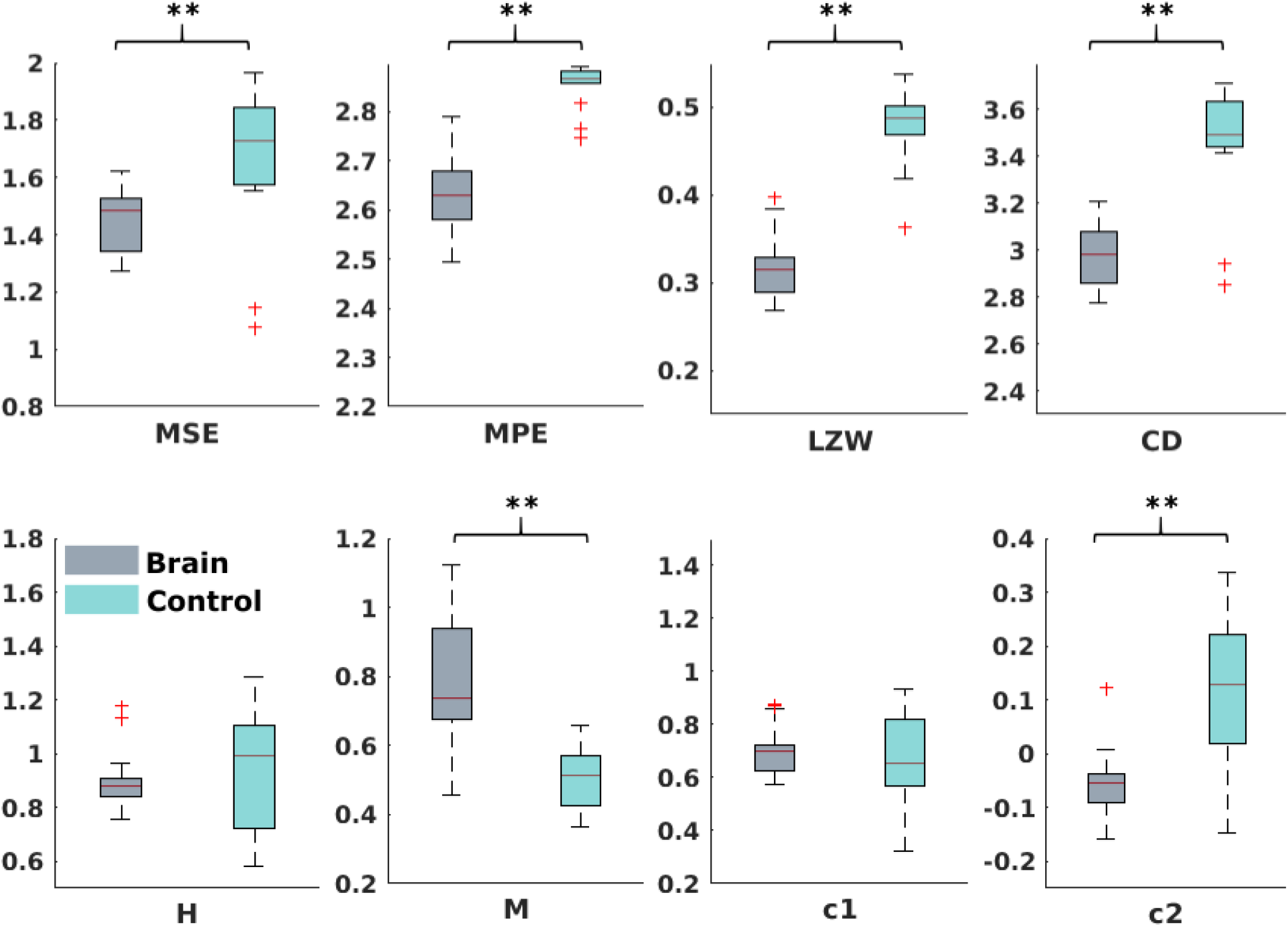
Results from analysis of different metrics on data from neonates without stimulation. ** depicts significant difference.

#### Fetal data

For the comparison of brain vs. control of in the fetal datasets, no clear tendencies could be found. The LZW calculation, as well as the MPE and CD metric did not reveal a significant difference, neither for audio data nor for spontaneous data. Some of the calculated metrics showed a difference for audio data only (MSE: p>0.001; H: p=0.02), one metric for both types of data (c1: p=0.01 audio; p=0.03 spont), and one metric for spont data only (c2: p=0.03). For detailed results, see Figures 4 and 5.

**Figure 4:**
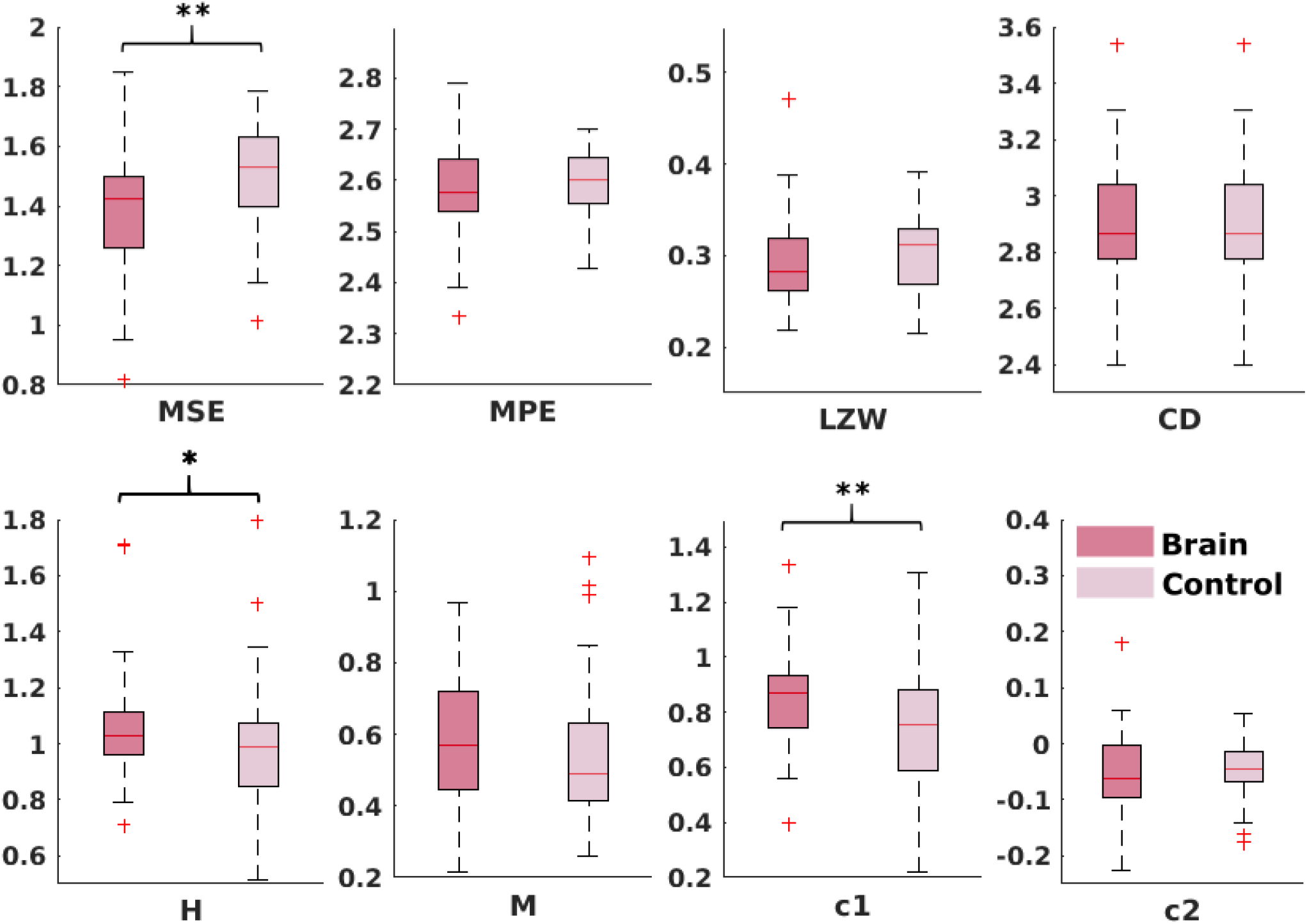
Results from analysis of different metrics on data from fetuses with auditory stimulation. * depicts significant, ** highly significant, difference.

**Figure 5:**
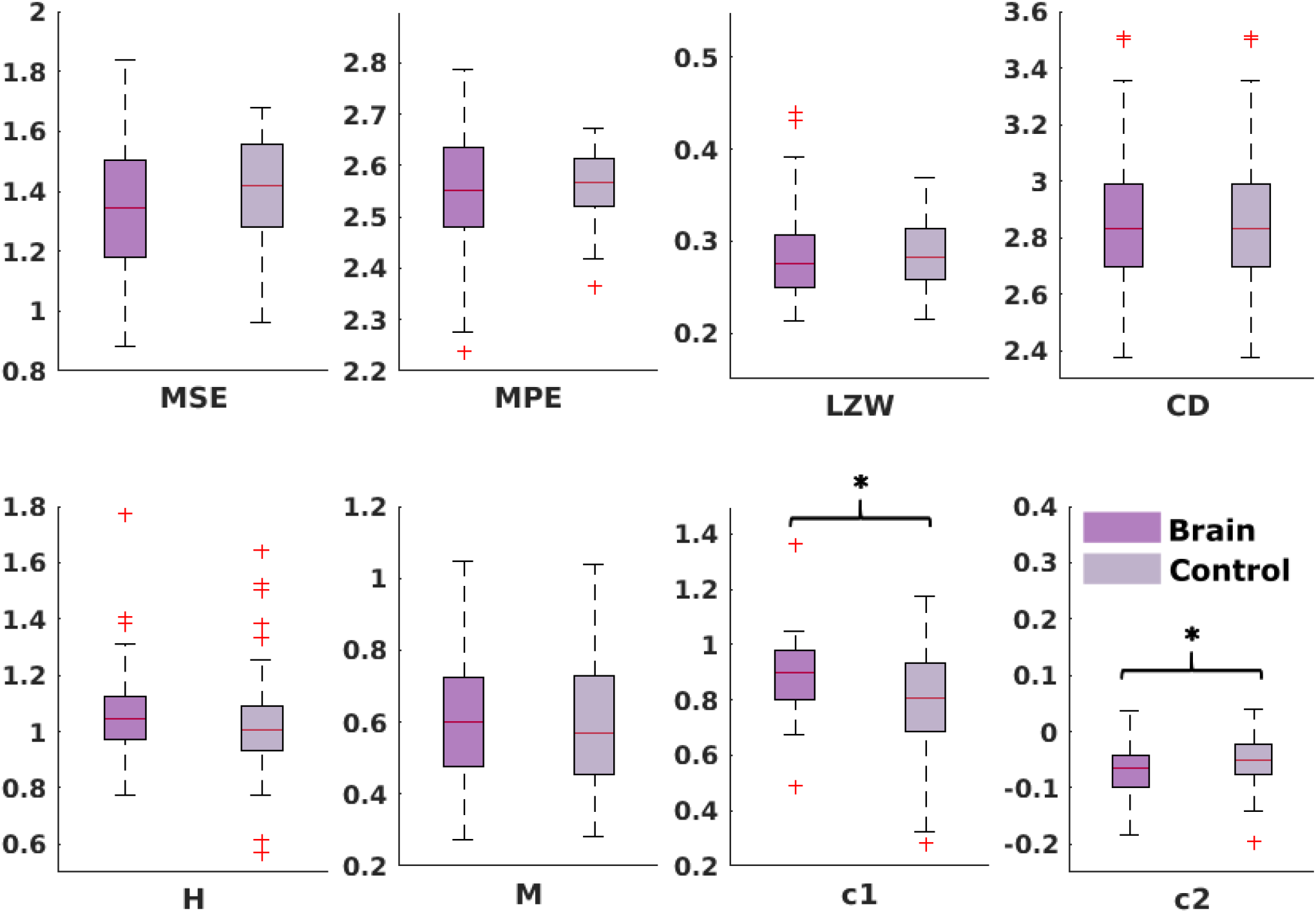
Results from analysis of different metrics on data from fetuses without stimulation. * depicts significant difference.

## 4 Discussion

The usability rating of the different metrics revealed that, concerning the ease of use with fMEG data, LZW was evaluated as best, as it is unequivocal and needs low computational effort. The fractality measures have a high parameter space and therefore forfeit comparability, while entropy measures require a higher computational effort and more parameters to adjust compared to LZW.

In the neonatal population, the channel with brain activity showed lower complexity compared to the control channel, measured by MSE, MPE, LZW and CD. As in this scenario, the control channel records environmental noise, it shows us that these metrics can clearly differentiate between a physiological signal and noise. These rather clear results can be seen as proof of concept for the general usability of complexity metrics in fMEG. For fetal data, this comparison produced less clear results – only MSE and scale free metrics showed a significant difference. This could be due to the fact that the control channel does not consist of pure environmental noise like in neonatal recordings but can also contain leftovers of physiological signals produced by fetus and mother. Magnetocardiographic activity of both, mother and fetus, should be taken into account as major confounds here. The different preprocessing steps that need to be performed, to evaluate fetal compared to neonatal data should be considered likewise.

The inconsistency of results from different metrics highlights the challenges of working with complexity metrics as neural correlates of consciousness, as well as the caution one should apply to interpret them. Especially, if a dataset consists of as many different aspects as fMEG data does, the choice of the right metric is crucial. Therefore, there is a need for more systematic, comparative studies, to evaluate the relations of different complexity metrics as well as their sensitivity to small changes in analysis parameters. Entropy measures during sleep for example, can reverse their direction, depending on the time scale used for calculation (Miskovic et al., 2019). Based on the assumption, that when using multiple scales to calculate entropy, random signals result in lower entropy values than complex signals (McIntosh et al., 2008), in the current study we would have expected higher MSE and MPE values for the brain compared to control channel, especially in neonatal data. Yet, we found the opposite in our results. This challenge especially accounts for metrics with high parameter space like the scale free metrics that even showed contradicting results in our analysis (M and c2 in neonatal data). The claim of these measures being equivalent (Zilber, 2014) did therefore not hold for the present results. Besides the analysis parameters, the preprocessing steps play a crucial role. We can see this for example in the relatively low LZW values in our results, that are related to the previous bandpass filtering of our data, as LZW values decrease with decreasing signal bandwidth (Aboy et al., 2006).

Further work is necessary to conclusively interpret results from this analysis of fetal MEG recordings. Even if largely explorative, this study shows that complexity metrics can be used for fMEG data on early consciousness and the evaluation gives a guidance for future work. The broad usage of LZW across a variety of studies, in combination with its use in earlier work on consciousness research, makes this metric especially interesting, as results can be compared to other subject populations. Yet, we need a better understanding of each metric and its sensitivity to different aspects of the data as well as their relation to different aspects of complexity, to use these empirical measures of complexity to assess the conscious state of a growing human being. However, as a precise assessment of fetal states is still challenging, the implementation of complexity metrics into fMEG research is a goal, that opens up interesting possibilities.

## 6 Author Contributions

JM conducted the analysis and wrote the manuscript. SB, EK revised the manuscript. JM, FS, HP conceptualized the study. SB, EK, GR, FW, HP advised the analysis. FS provided preceding analysis. All authors read and approved the final version of the manuscript.

## 8 Acknowledgments

This work was presented as a poster at the HBP International Conference “Understanding Consciousness; A scientific quest for the 21^st^ century”, June 21^st^-22^nd^, 2018 in Barcelona.

## Footnotes

1 Multiscale permutation entropy (MPE), version 1.3. Retrieved from https://ch.mathworks.com/matlabcentral/fileexchange/37288-multiscale-permutation-entropy–mpe

2 Chaotic systems toolbox, version 1.0. Retrieved from https://ch.mathworks.com/matlabcentral/fileexchange/1597-chaotic-systems-toolbox

3 Minimum embedding dimension, version 1.0. Retrieved from https://ch.mathworks.com/matlabcentral/fileexchange/37239-minimum-embedding-dimension

